# Partitioned Multi-MUM finding for scalable pangenomics

**DOI:** 10.1101/2025.05.20.654611

**Authors:** Vikram S. Shivakumar, Ben Langmead

## Abstract

Pangenome collections are growing to hundreds of high-quality genomes. This necessitates scalable methods for constructing pangenome alignments that can incorporate newly-sequenced assemblies. We previously developed Mumemto, which computes maximal unique matches (multi-MUMs) across pangenomes using compressed indexing. In this work, we extend Mumemto by introducing two new partitioning and merging strategies. Both strategies enable highly parallel, memory efficient, and updateable computation of multi-MUMs. One of the strategies, called string-based merging, is also capable of conducting the merges in a way that follows the shape of a phylogenetic tree, naturally yielding the multi-MUM for the tree’s internal nodes as well as the root. With these strategies, Mumemto now scales to 474 human haplotypes, the only multi-MUM method able to do so. It also introduces a time-memory tradeoff that allows Mumemto to be tailored to more scenarios, including in resource-limited settings.

## 1 Introduction

Pangenome assembly collections continue to proliferate and grow. Recently published pangenomes have covered crops (Garg et al. 2024), model organisms (Hufford et al. 2021; Lian et al. 2024), and the expanded v2 version of the Human Pangenome Reference Consortium (HPRC) dataset spanning 474 human haplotypes (Liao et al. 2023; H Li 2025). Other recent pangenomes span distinct but related species (Yoo et al. 2025), as well as geographically and genetically distinct accessions of the same species (Lian et al. 2024). Because of the growing size and complexity of these collections, it is increasingly important that our methods for studying and indexing pangenomes be scalable. Specifically, we wish to be able to find conserved elements across pangenomes in a way that scales to large pangenomes and that can be updated incrementally when the pangenome grows further. Such methods will allow us build the common coordinate systems that will underlie future studies (Taylor et al. 2024).

A multiple Maximal Unique Match (multi-MUM) is an exact sequence match that is present exactly once in all the pangenome sequences. Because they are both unique in each sequence and common to all the sequences, multi-MUMs serve as particularly strong guideposts for establishing common pangenome coordinates. They have been used as anchors for multiple alignment (Kille et al. 2024; Deogun et al. 2004; Darling et al. 2010; Olbrich et al. 2025) and as markers for chaining of pangenome read alignments (Brown et al. 2025). We previously proposed a framework and tool called Mumemto for rapidly identifying multi-MUMs during construction of a compressed pangenome index (Shivakumar Langmead 2025). Using prefix-free parsing (PFP) (Boucher et al. 2019), a compressed-space method for computing full-text indexes, Mumemto outperforms existing methods for identifying sequence-level multi-MUMs. Another method, PANAMA, uses a similar PFP-based approach to compute larger syntenic blocks for multiple alignment. However, while both of these tools were capable of operating over pangenomes of moderate size, PFP cannot currently be scaled to the larger collections available today. For instance, running Mumemto on the 474 human haplotypes of the HPRC v2 release (Liao et al. 2023; H Li 2025) requires over 2TB of memory.

To address this, we developed a partition-merging approach to compute multi-MUMs with Mumemto. The method separates the input genomes into partitions, then uses PFP to compute per-partition intermediate results, comprising multi-MUMs and additional match information. The method then merges results from the partitions to yield a global set of multi-MUMs across the union of the partitions. While past methods like Parsnp have proposed ways to iteratively merge pairwise MUMs into multi-MUMs (Treangen 2008; Treangen et al. 2014), our method generalizes to merging across arbitrary initial collections of sequences, including multi-way merging. This yields a highly parallelizable and easily updateable multi-MUM-finding workflow that scales to hundreds of human genomes.

We propose two methods for merging multi-MUMs. “Anchor-based merging” uses a common reference sequence in each partition to anchor matches and identify overlaps between subsets for subsequent merging. “String-based merging” finds overlaps between multi-MUMs from the sequence directly, merging multi-MUMs across disjoint sequence collections. When applying string-based merging to partitions that correspond to evolutionarily related clades, the intermediate results computed at each step are the multi-MUMs for some internal node of the phylogenetic tree. As a result, we can establish a set of multi-MUMs, and therefore a common coordinate system, at multiple levels of the phylogeny at once.

By implementing a partition-merging scheme for computing multi-MUMs, we also introduce a tradeoff space between highly-parallelizable multi-MUM computation and peak memory usage. This allows Mumemto to scale to hundreds of human genomes by splitting the genomes into partitions and processing each partition serially. It also allows Mumemto to accelerate multi-MUM finding by processing the partitions in parallel, at the expense of a higher total memory footprint. Given this merging strategy, Mumemto can also now easily integrate newly-assembled genomes without recomputing from scratch, making it well-suited to scale as pangenome collections continue to grow in the future and as a core algorithm for pangenome construction and analysis.

## 2 Methods

### 2.1 Preliminaries

*Consider text T*, a concatenation of *N* sequences [*T*_0_ … *T*_*N*_], with total length *n* over alphabet Σ. The *i*^th^ suffix is defined as the substring *T* [*i*..*n*]. The suffix array *SA* of *T* contains the offsets [0..*n*] of *T*, ordered by lexicographic rank of the corresponding suffix. The Burrows-Wheeler Transform *BWT*_*T*_ is a sequence consisting of characters that precede each of *T* ‘s suffixes in the order given by *SA*, i.e. *BWT*_*T*_ [*i*] = *T* [*SA*[*i*]−1] (assuming the text is circular). The longest common prefix array at offset *i, LCP* [*i*], contains the longest common prefix between suffixes *T* [*SA*[*i* − 1]..*n*] and *T* [*SA*[*i*]..*n*] (with *LCP* [*i*] = 0). The document array *DA* contains the text of origin for each suffix in suffix array order, e.g. for *DA*[*i*] = *k, T* [*SA*[*i*]..*n*] begins in *T*_*k*_ in the concatenation of sequences. In practice, text *T* contains a delimiter between each sequence that is lexicographically smaller than any character in Σ.

### 2.2 Mumemto

Mumemto (Shivakumar Langmead 2025) computes various types of exact matches. These might be constrained to be unique in each sequence (i.e. a Maximal Unique Match or MUM), or not (i.e. a Maximal Exact Match or MEM). They might also be required to appear in all sequences (a multi-MUM or multi-MEM), or only in some fraction of them (a partial multi-MUM or partial multi-MEM). In this work, we focus specifically on multi-MUMs. We may refer to them as simply MUMs for short when the “multi” is clear from context. Multi-MUMs are represented by an interval [*i, j*] of size *N* in the *SA, BWT, LCP* array, and *DA*. This window has the following properties:

1. **(exact match)** With *l* = min *LCP* [*i* + 1..*j*] representing the length of the shared common prefix among suffixes in the interval, *LCP* [*i* + 1] *> l > LCP* [*i*] and *LCP* [*j*] *> l > LCP* [*j* + 1].
2. **(uniqueness)** *DA*[*i*..*j*] contains pairwise distinct values
3. **(left-maximal)** the characters in *BWT* [*i*..*j*] are not all identical

Mumemto adapts an algorithm ((Shivakumar Langmead 2025), Algorithm 1) proposed by Abouelhoda et al. (Abouelhoda et al. 2004) to find multi-MUMs by identifying intervals in the various arrays where these properties hold.

### 2.3 Prefix-free Parsing

Prefix-free parsing (PFP) (Boucher et al. 2019) is a compressed-space algorithm for computing the BWT, SA, and LCP array. It begins by parsing the input text into distinct phrases (the dictionary *D*) and storing the sequence of phrases that make up the original text (the parse *P*). By computing the SA, LCP, and BWT over both *D* and *P*, PFP can reconstruct the original arrays, requiring only *O*(|*D*| + |*P*|) space. The space requirement tends to grow sublinearly for repetitive texts and represents the amount of unique sequence present in the pangenome (Lipták et al. 2025). Crucially, PFP computes and reports these arrays in order and in tandem, allowing Mumemto to scan for windows satisfying the properties mentioned above (exact match, uniqueness, left-maximality) in a streaming fashion and without writing the arrays to disk.

### 2.4 Anchor-based merging

We propose a pair of related algorithms for merging multi-MUMs computed across *k* pangenome partitions. In both cases, the final set of merged multi-MUMs is the same as if they had been computed over the union of the merged partitions. We propose two methods: one that uses the coordinate system of a designated pangenome representative to “anchor” the merging process (“anchor-based merging”), and a second that does not require a common sequence across partitions and merges purely based on the sequences of the multi-MUMs themselves (“string-based merging”). The choice of anchor sequence is arbitrary, due to the fact that a multi-MUM appears in all sequences by definition, and so appears at some offset in any particular sequence.

Merging multi-MUMs across partitions requires some care. A simple case is one where the two partitions each have a corresponding multi-MUM that, once the partitions are merged, is still a multi-MUM in the merged collection. But subtler cases arise when (a) a multi-MUM in one partition is no longer a multi-MUM in the merged collection due to genetic variation or differences in repetitive sequence, or (b) multi-MUMs in the different partitions combine to form a merged multi-MUM that is shorter than one or both of the originals, due to genetic differences that are unique to one partition or the other. To handle all such cases, we store extra information for each partition that indicates how short a match can be truncated before it is no longer unique in the given partition. To do so, we consider unique matches (UMs), which contain properties (1) and (2) in section 2.2, but not property (3), i.e. they may be further extendable to the left. While traversing the arrays, for every multi-MUM or UM encountered, we store max(*LCP* [*i*], *LCP* [*j* + 1]), which represents the longest prefix of the MUM or UM that appears elsewhere in the text. In other words, if the MUM or UM is truncated below this length from the right, it will no longer be unique in at least one sequence in the partition. We store this length at the offset where the MUM or UM occurs in the anchor reference sequence.

We store such a value for each offset of the anchor reference, yielding an array called `unique_threshold`. Each value in `unique_threshold` is either 0, indicating no UM or MUM starts at that offset, or *l >* 0, indicating the match occurring at that offset will become non-unique if truncated to a length shorter than *l*. Algorithm 1 describes and Figure 1 illustrates merging two partitions using this array. This can be generalized to merging *k* partitions by outputting an updated `unique_threshold` array for the merged output, which can be used in a subsequent round of merging.

**Figure 1:**
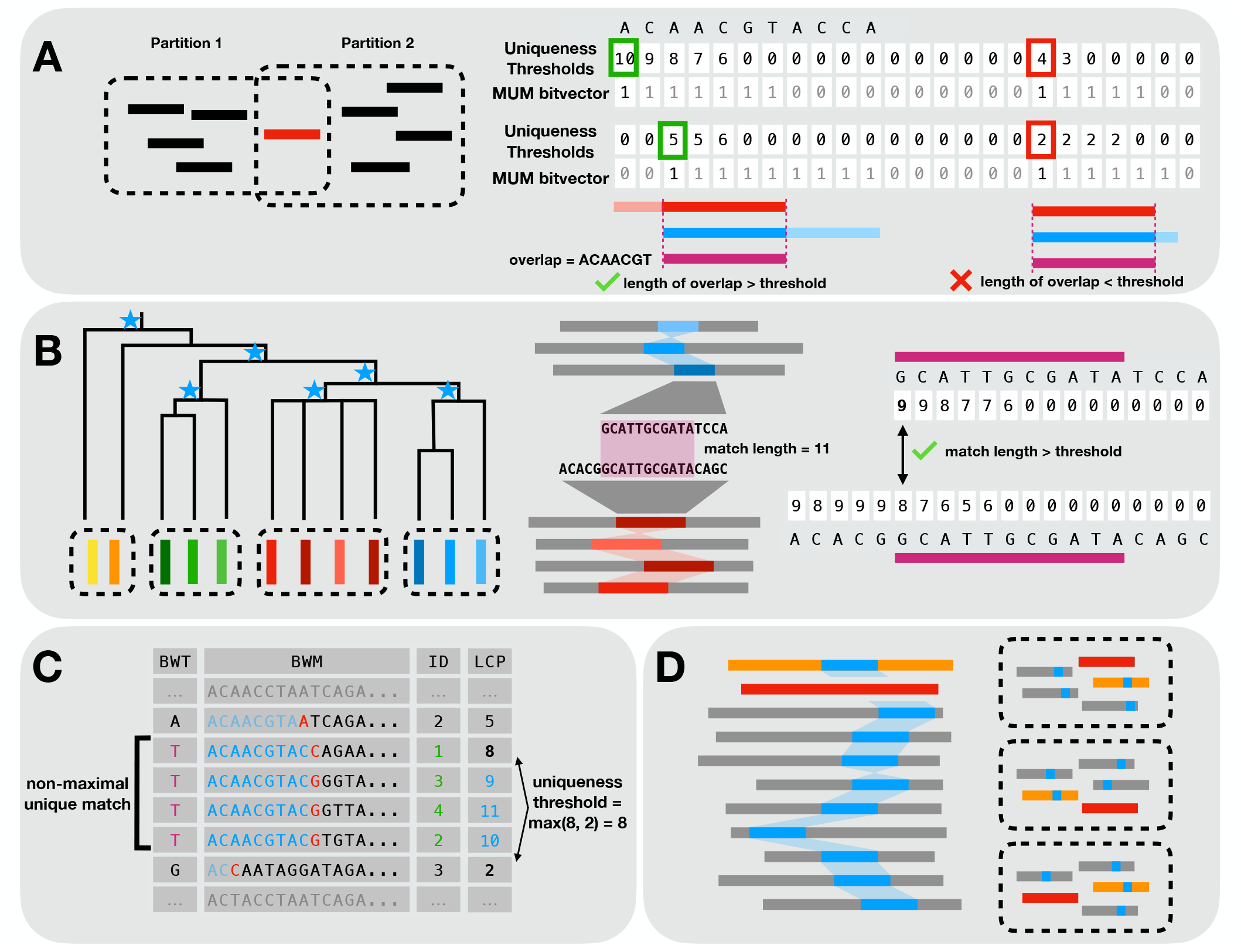
(**A**) Anchor-based merging requires a common sequence (red) present in each partition. Multi-MUMs are merged by identifying overlaps between partition-specific matches in the anchor coordinate space, and a uniqueness threshold determines if a MUM is still unique in each partition after truncation. (**B**) String-based merging enables computation of multi-MUMs between partitions without a common sequence. An example tree (left) is shown, highlighting the use case where partial multi-MUMs specific to internal nodes (starred) can be computed by merging subclade-based partitions up a tree. (right) MUM overlaps are computed by running Mumemto on the MUM sequences, and the uniqueness threshold array ensures overlaps remain unique across the merged dataset. (**C**) An example Burrows-Wheeler Transform (BWT), matrix (BWM), and Longest Common Prefix (LCP) array, with sequence IDs for each suffix shown (ID). A non-maximal unique match (UM) is shown, and the uniqueness threshold for this match is found using the flanking LCP values. (**D**) A partial multi-MUM (in blue) is found in all-but-one sequence (excluded in red). Using two anchor sequences (red and orange), all-but-one partial MUMs can be computed using an augmented anchor-based merging method (section 2.6).

#### Algorithm 1 Anchor-based MUM Merging

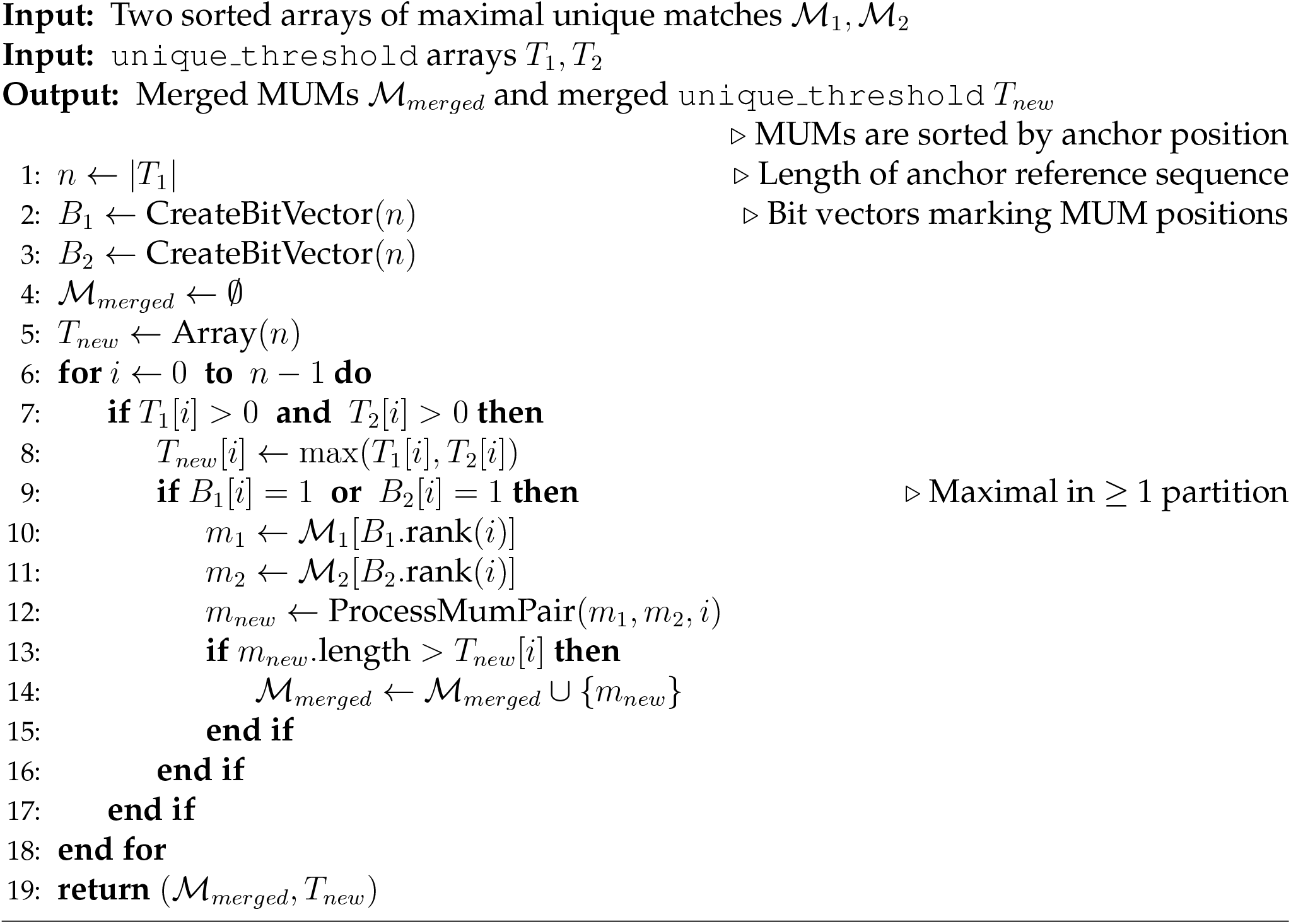

### 2.5 String-based merging

The anchor-based merging strategy requires a common reference sequence across partitions. We now propose a method for multi-MUM merging that works for disjoint partitions with no sequences in common. Because this strategy uses the sequences of the multi-MUMs themselves as the basis for its merging decisions, we call it the string-based merging strategy.

We again store the `unique_threshold` array; as before, this array indicates how short a MUM or UM can be truncated before it loses uniqueness in a partition. However, we now store these values for each offset of each MUM yielding an array of size *O*(total length of MUMs), which is usually shorter than the length of the anchor reference. Additionally, we store this information for the reverse complement of each MUM or UM. This is used to construct the merged `unique_threshold` array for subsequent merging cycles.

To identify overlaps, we extract multi-MUM sequences from each of the *k* partitions being merged and re-invoke Mumemto on those. That is, the collection of multi-MUMs from each partition becomes the new “genome” and we now find its multi-MUMs. When we detect an overlap between matches present in all *k* partitions, we check the `unique_threshold` arrays for each to determine whether the multi-MUM should be kept as-is, truncated, or removed, similarly to the anchor-based method. Lastly, we build a new `unique_threshold` for the merged dataset by computing an element-wise comparison over threshold values from the overlap regions across partitions. We use the reverse-complement thresholds when the overlap is on the negative strand of a MUM. Algorithm 2 details this procedure.

#### Algorithm 2 String-based MUM Merging

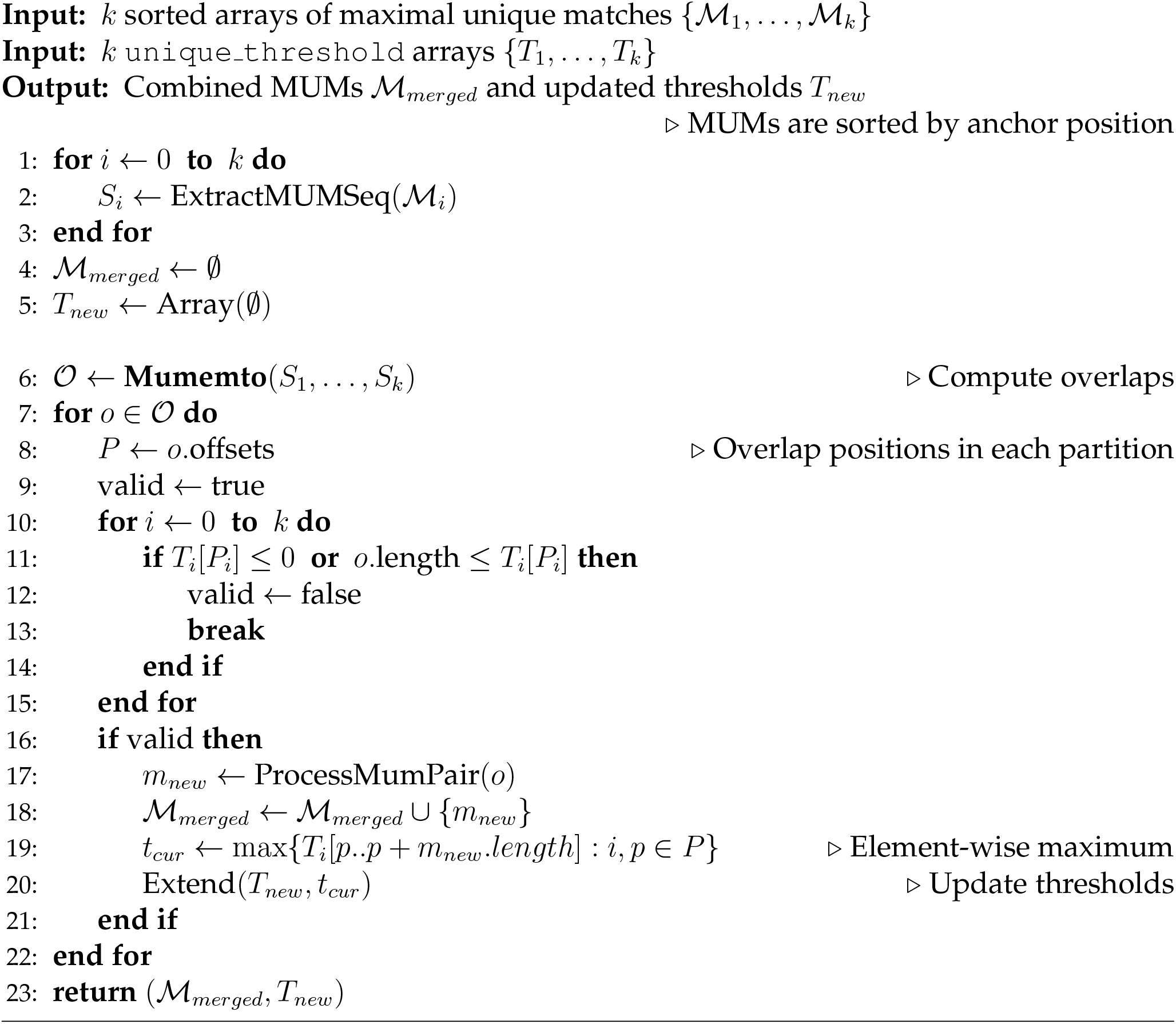

### 2.6 Extension to partial multi-MUMs

The method described so far works exclusively for multi-MUMs. In previous work (Shivakumar Langmead 2025), we showed the utility of reporting other match types, e.g. matches that appear in a large subset of the genomes but not all. We now describe an extension to the anchor-based merging method that enables computation of partial multi-MUMs appearing in *N* − 1 sequences (all-but-one), as well as ideas for generalizing to the case of artial multi-MUMs appearing in at least *N* − *p* sequences for small *p >* 1.

To find partial multi-MUMs present in *N* −1 sequences, we must handle the case where a multi-MUM appears in all sequences except the anchor reference sequence. Anchor-based merging fails in this case since the MUM cannot be assigned an offset with respect to the anchor. To address this, we instead select *two* anchor sequences, one that serves as the primary anchor sequence for most partial MUMs, and a second that serves as a “backup” anchor sequence for the partial MUMs that do not appear in the primary anchor sequence. Additionally, we store a `unique_threshold` *for each anchor reference*. We compute partial multi-MUMs appearing in *N* − 1 sequences for each partition (i.e. Mumemto flag `-k -1`). If a match appears in the first anchor reference, we store the `unique_threshold` at the first anchor offset. If a match does not appear in the first anchor sequence (which implies it must appear in all other sequences), we store the threshold information at the second anchor offset. We proceed with merging as normal, comparing the first and second anchor thresholds across partitions. However, we only process overlaps that are partial in exactly one partition, ensuring the final merged MUM is appears in at least *N* − 1 sequences.

To generalize to partial multi-MUMs appearing in at least *N* − *p* sequences for *p >* 1, we must now select a set of *p* + 1 common anchor sequences, and include all of them in all partitions. Assume that the anchor sequences have a priority order, with one being primary, another secondary, etc. Then each of these anchor references serves as a “backup” for the case of a partial multi-MUM that fails to appear in any of the higher-priority anchor references. Since the addition of each new anchor sequence adds overhead to the initial partition-wise MUM-finding step, this becomes impractical unless *p* is small.

## 3 Results

We implemented both anchor and string-based merging in Mumemto for computing multi-MUMs. Users can enable mergeability by running Mumemto with the (-M) option, which causes Mumemto to report the auxiliary `unique_threshold` array in addition to multi-MUMs. For anchor-based merging, this array uses 2*n* bytes, where *n* is the length of the anchor reference sequence. For string-based merging, it uses 4*m* bytes, where *m* is the total length of multi-MUMs.

To benchmark and apply the paritition-merging method, we used assemblies from the Human Pangenome Reference Consortium (HPRC) (Liao et al. 2023), specifically 474 haplotype assemblies (available from (H Li 2025)). To obtain chromosome-level assemblies for each haplotype, we performed reference-guided scaffolding against CHM13 using RagTag (default parameters) (Alonge et al. 2022). We only considered contigs which were included in the scaffolded assembly when referring to HPRC assemblies below.

### 3.1 Time and memory tradeoffs

We implemented the methods of section 2 in a workflow that (a) breaks the pangenome into partitions, (b) computes multi-UMs in each partition, then (c) merges the per-partition matches into a global set of multi-MUMs. The user can trade between running time and peak memory usage by setting partition size and number of threads. Running parallel, per-partition Mumemto processes and merging the results reduces the running time but increases the peak memory footprint. Running a single Mumemto thread over each partition serially yields longer running time but a smaller memory footprint.

We measured the time and memory efficiency of running Mumemto over a partitioned dataset in parallel and serially for two datasets using anchor-based merging. The datasets were human (Liao et al. 2023) and *A. thaliana* (Lian et al. 2024). Splitting 474 chr19 haplotypes into 40 partitions and processing those fully in parallel yielded an 18.3 × speedup (0.26 hours versus 4.77 hours for 1 thread) but used 4.3 × more memory (190.39 GB versus 44.05 GB for 1 thread). When partitioned and processed serially, the memory usage was reduced by 8.2 × (5.35GB versus 44.05GB for 1 thread), with a 1.7 × increase in runtime (from 4.77 hrs to 8.32). For *A. thaliana*, processing the 69 accession assemblies in 20 partitions in parallel yielded a 6.5 × speedup (0.40 hours versus 2.58 hours for 1 thread) with a 3.4 × memory increase (242.36 GB versus 70.34GB for 1 thread). Run serially, a 5 × memory reduction (from 70.34GB to 13.58GB) is achieved with a 2.5 × increase in runtime (from 2.58 hrs to 6.48).

The *A. thaliana* tradeoff is similar though smaller in magnitude compared to human assemblies. Recall that the anchor-based merging method introduces overhead by including the common anchor reference in each partition. When the sequences being indexed are highly similar, e.g. human genome assemblies, then PFP compression can effectively mask this additional overhead. But for the more divergent *A. thaliana* dataset, the additional overhead is more apparent, leading to the narrower trade-off space.

**Table 1:** Runtime and memory usage comparison between serial and parallel multi-MUM computation across different partition schemes. We report execution time (in hours) and peak memory usage (in GB) for two datasets, chr19 haplotypes from the Human Pangenome Reference Consortium (HPRC) (Liao et al. 2023) and the *A. thaliana* pangenome (Lian et al. 2024). The lowest peak memory footprint and the fastest running times are bolded.

| Dataset | Partitions | Seqs per | Serial |  | Parallel |  |
| --- | --- | --- | --- | --- | --- | --- |
|  |  |  | Time (hrs) | Memory (GB) | Time (hrs) | Memory (GB) |
| chr19 HPRC<br>($N = 474$) | 1 | 474 | 4.77 | 44.05 | – | – |
|  | 5 | 96 | 5.00 | 13.96 | 1.11 | 66.78 |
|  | 10 | 48 | 5.91 | 9.56 | 0.63 | 88.63 |
|  | 20 | 24 | 7.21 | 7.16 | 0.39 | 125.29 |
|  | 40 | 12 | 8.32 | **5.35** | **0.26** | 190.39 |
| *A. thaliana*<br>($N = 69$) | 1 | 69 | 2.58 | 70.34 | – | – |
|  | 5 | 15 | 3.75 | 27.14 | 0.80 | 128.08 |
|  | 10 | 7 | 4.92 | 18.79 | 0.57 | 172.00 |
|  | 20 | 4 | 6.48 | **13.58** | **0.40** | 242.36 |

### 3.2 Anchor-based merging across 474 human haplotypes

The Human Pangenome Reference Consortium (Liao et al. 2023) released 474 human haplotype assemblies (available at (H Li 2025)), totaling ∼1.4 Tbp (or 2.8 Tbp with reverse complements). Running Mumemto over the entire collection, even with PFP compression, would be infeasible, with a total PFP size of 145.6 GB, requiring over 2.7 TB of memory. However, we note that the primary bottleneck in this case is the size of the parse rather than the dictionary, which recursive extensions to PFP aim to address (Ferro et al. 2024).

With CHM13 as the common coordinate system for merging, we divided the HPRC dataset into 9 partitions (53–54 assemblies in each) and computed multi-MUMs within each subset. We note that although the choice of anchor reference does not affect the final multi-MUM output, CHM13 provides a convenient choice as the most complete T2T assembly for humans. Merging across subsets, we found 20,914,049 multi-MUMs across all 474 assemblies, totaling ∼2Gbp of non-overlapping sequence in CHM13 and an average coverage of 73.2% across haplotypes. The updated HPRC v2 dataset multi-MUM coverage is smaller than that of the previously released v1 (89 haplotypes) (84.8% coverage, totaling 2.43Gbp of sequence). This is expected, as the dataset grows, deletions and incomplete assemblies in newly added haplotypes will lower overall multi-MUM coverage. However, including partial MUMs in all-but-one sequences in HPRC v1 increases MUM coverage to 86.2%, motivating the merging algorithm extension for partial MUMs detailed in section 2.6.

We combined collinear MUMs separated by *<* 100bp into “blocks,” as described in previous work (Shivakumar Langmead 2025). We found 254,440 collinear MUM blocks, covering 76.4% of CHM13. A collinear block contained 82 multi-MUMs on average, with over two-thirds of the gaps between collinear multi-MUMs being a single base wide (consistent with previous work (Shivakumar Langmead 2025)). Run serially, the partitioned Mumemto pipeline would take 339.33 hrs (a little over 2 weeks), using 448.20 GB of memory. Using our partitioning and merging strategy to run this same process across 9 compute nodes in parallel, the pipeline took 40.36 hrs using roughly 434 GB of memory per node. Mumemto is the only method that can currently compute multi-sequence exact matches across this full set of 474 haplotypes. Further, Mumemto’s merging strategy allows for incorporation of newly-released human assemblies, without needing to recompute multi-MUMs over the assemblies indexed previously.

### 3.3 Merging multi-MUMs across a phylogenetic tree

In cases where a pangenome spans individuals of multiple species, or spans multiple diverse accessions, the sequences’ natural groupings may be determined by a tree. For instance, the recent *A. thaliana* pangenome (Lian et al. 2024) spans genomes from geographically diverse accessions, which can be related in a phylogenetic tree. In this case, we can use string-based merging to allow the partitioning and merging process to follow the structure of tree bottom-up, so that the intermediate results correspond to the tree’s internal nodes (Figure 1B). This is possible because string-based merging allows sibling subtrees to be merged without a common anchor reference. An additional benefit is that the intermediate computations will always be over clades of the tree, which consist of more similar genome sets than if we had partitioned arbitrarily. This maximizes PFP compression in each subproblem.

We applied this strategy to a phylogeny and collection of *A. thaliana* assemblies from diverse geographical regions (Lian et al. 2024) (Figure 2A). Figure 2B shows the multi-MUM synteny across the full dataset. Multi-MUM coverage across the full dataset drops in the highly-variable centromeric region, however, much of the synteny is revealed by local partition-specific MUMs. Highlighting these partition-specific MUMs reveals novel structural variation local to geographically isolated subgroups. For instance, a 778 Kbp inversion is present in the chr4 centromere of two Madeiran accessions (Figure 2D), and a complex inversion and rearrangement region is revealed by partial MUMs among accessions from North America in the chr5 centromere (Figure 2E).

**Figure 2:**
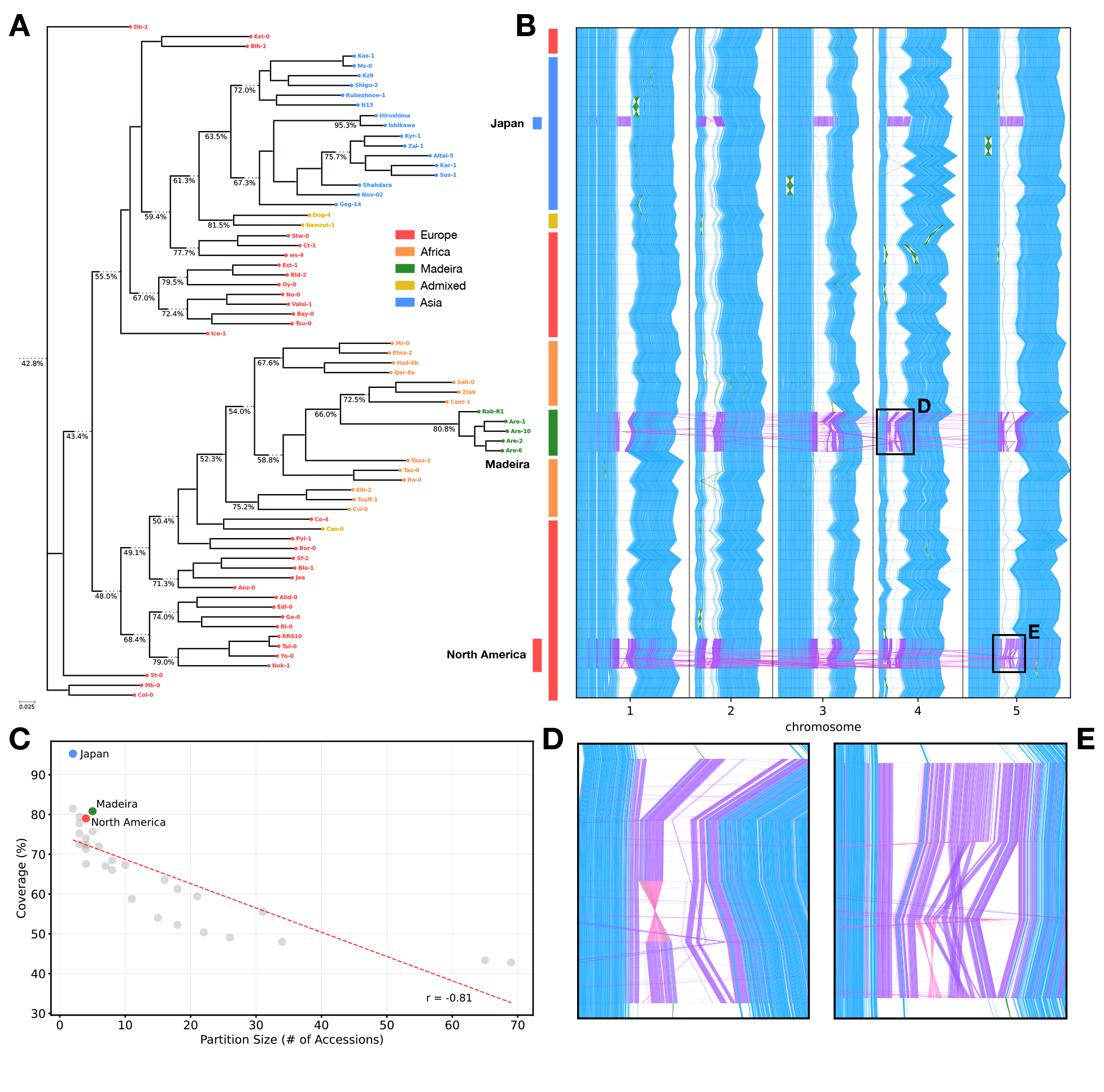
(**A**) Phylogeny of geographically diverse *A. thaliana* accessions (Lian et al. 2024), with broad geographical regions colored. Internal nodes are labeled with the coverage of partial multi-MUMs across the leaves of each node. Internal node partial MUMs are computed by merging subtree-based partitions progressively up the phylogeny. (**B**) Global multi-MUM synteny across the full dataset shown in blue (with inversions in green). Global MUMs are computed by merging all partitions together (representing the root node). Additionally, three geographically distinct subgroups are highlighted and partition-specific multi-MUMs (in purple, with inversions in pink) reveal local structural variation in centromeric regions. Two examples are shown in zoomed-in panels: (**D**) a novel inversion located in accessions from the Madeira islands, and (**E**) a complex inversion/rearrangement in the North American subgroup chr5 centromere. (**C**) Coverage of internal nodes in the phylogeny compared to number of leaves under node. Three highlighted subgroups show particularly high coverage, likely due to geographical isolation (Lian et al. 2024).

Across the full dataset, multi-MUM coverage is 42.8%, which corresponds well with previously reported measures of collinearity (Lian et al. 2024). As expected, MUM coverage drops as partitions are merged up the tree (Figure 2A). Lian et al. also reported increased collinearity in certain small geographical groups. Similarly, we found an increased multi-MUM coverage of 95% in the Japanese accessions, 79% in the North American subgroup, and 81% in the Madeiran subgroup (Figure 2C). By computing multi-MUMs in a hierarchical manner, we can find subgroup-specific conserved regions and dataset-wide multi-MUMs with a single, partitioned Mumemto run.

### 3.4 Merging enables inter-specific multi-MUM computation

PFP is designed for repetitive inputs like pangenomes consisting of many same-species genomes. Inter-specific datasets have greater sequence diversity and are less compressible. Consistent with this, we observed that the prefix-free parse of 28 primate genome assemblies (from a curation by Hecker and Hiller, 2020 (Hecker Hiller 2020) and the T2T primate project (Yoo et al. 2025)) was larger than the original sequence files. As a result, computing multi-MUMs across the 28 primate assemblies (∼150Gbp) with non-partitioned Mumemto required over 2TB of memory. Using partitioned Mumemto, however, we can reduce the memory required for computing multi-MUMs across diverse, inter-specific datasets like this one. By partitioning the genomes into 4 groups and processing the partitions serially, we were able to find multi-MUMs across the primates in 842 GB of memory and 37.4hrs of running time.

To further explore this new multi-MUM dataset, we studied the most conserved sequences across the primate species. Similar to ultraconserved elements (UCEs) (Hecker Hiller 2020; Bejerano et al. 2004; Cummins et al. 2024), multi-MUMs across diverse genomes can be used to identify regions of evolutionary conservation. We identified collinear blocks from multi-MUMs present across 29 primate assemblies (including GRCh38 and CHM13). We considered all collinear blocks (with maximum space between MUMs of 100bp) and non-collinear MUMs longer than 50bp to be conserved elements across primates. We found 10,382 such elements, totaling ∼1.56Mbp of sequence. The average element length was 150bp, with 198 elements longer than 500bp. Most of these conserved sequences were located in non-protein-coding regions (83.81%), and roughly half (42.13%) in intergenic regions (using MANE 1.4 to define genic and protein-coding regions (Morales et al. 2022)). We also found 1,410 (13.6%) elements spanned intron-exon boundaries. These results are consistent with previously reported genomic distributions of ultraconserved elements in humans (Cummins et al. 2024), highlighting the potential for inter-specific multi-MUMs for identifying UCEs.

## 4 Discussion

Multi-MUM merging algorithms enable a partition-based, memory-efficient, and parallelizable Mumemto workflow, which now scales to hundreds of human genomes. These algorithms also simplify incorporating newly assembled genomes by dynamically updating multi-MUMs through merging, rather than recomputing MUMs across the full, updated dataset. This combination of scalability and updatability makes Mumemto well suited to a future where long-read sequencing projects produce steady streams of new assemblies.

The two merging schemes discussed each have their own limitations. Anchor-based merging is faster because finding multi-MUM overlaps is easily done with numerical comparisons in a common coordinate system. On the other hand, this requires that a common anchor reference be present across the partitions, adding both time and space overhead. This is still suitable when the goal is to simply compute multi-MUMs over a large dataset such as HPRC v2. String-based merging is slower than anchor-based merging, since it requires an additional invocation of Mumemto to identify overlaps. However, it enables tree-based merging to find partial multi-MUMs at internal nodes, revealing conservation and variation between subgroups in the dataset.

String-based merging relies on running Mumemto over the MUM sequences themselves to identify overlaps. This enables merging multiple partitions at once, although we find that memory usage of the merging step can become the limiting factor for a large number of partitions. We note that any exact match method could be used to find these overlaps (for example, Mummer4 (Marçais et al. 2018)). We used Mumemto itself, in order to limit external dependencies. In the future it will be important to investigate if another exact-matching method can accelerate string-based merging and close the speed gap with anchor-based merging.

Both merging methods are limited to computing strict multi-MUMs, however we described an extension that enables partial multi-MUM finding in all-but-a-few sequences. A question for future work is whether a more sophisticated method could enable computing general partial multi-MUMs across any subsets.

We show that a partitioned Mumemto enables scalability to growing pangenome collections and expands the applicability of Mumemto to larger, more diverse datasets. This increases the scope for exploration of genomic conservation and variation and highlights the potential for Mumemto as a core method for future pangenomics and comparative genomics research.

## 5 Data Availability

The methods presented here are implemented in the Mumemto software (v1.3.0), available open-source at https://github.com/vikshiv/mumemto. Mumemto v1.3.0 was used for all analyses.

## 7 Acknowledgements

The authors would like to thank Christina Boucher and Travis Gagie for algorithmic discussions, and Kuan-Hao Chao for valuable discussions on human genome annotations.

## 8 Authors’ Contributions

V.S.S. and B.L. conceived the project. V.S.S. wrote the software and conducted the experiments. V.S.S. and B.L. wrote the manuscript.

